# BioBlocks: Programming protocols in biology made easier

**DOI:** 10.1101/081075

**Authors:** Vishal Gupta, Jesús Irimia, Iván Pau, Alfonso Rodríguez-Patón

## Abstract

The methods to execute biological experiments are evolving. Affordable fluid handling robots and on-demand biology enterprises are making automating entire experiments a reality. Automation offers the benefit of high-throughput experimentation, rapid prototyping and improved reproducibility of results. However, learning to automate and codify experiments is a difficult task as it requires programming expertise. Here, we present a web-based visual development environment called BioBlocks for describing experimental protocols in biology. It is based on Google’s Blockly and Scratch, and requires little or no experience in computer programming to automate the execution of experiments. The experiments can be specified, saved, modified and shared between multiple users in an easy manner. BioBlocks is open-source and can be customized to execute protocols on local robotic platforms or remotely i.e. in the cloud. It aims to serve as a ‘de facto’ open standard for programming protocols in Biology.

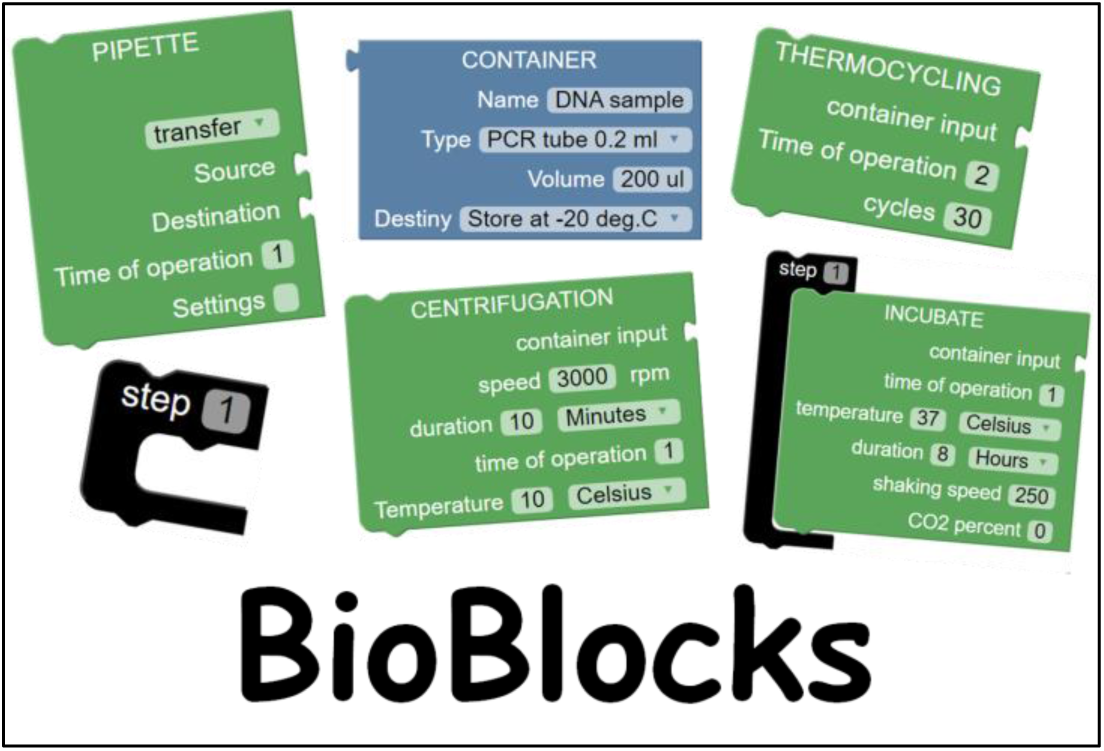

## 1. INTRODUCTION

The inability to reproduce the results of biological research has become a critical issue to address because of its economic and scientific impact.^1^ Some factors that contribute to the problem of reproducibility are the ambiguity introduced by natural languages (English) when describing experiments, the person-to-person variability while carrying out experiments, inadequate data sharing, poor quality control, etc. Several academic and commercial solutions address these problems using automation. Biological protocols described using programming languages are precise and can be automated as the description (code) is machine-readable. However, they have not been successful because it requires the user (biologist) to have expertise in programming. Notable efforts in this direction are academic solutions like BioCoder^2^, Puppeteer^3^, AquaCore^4^, Par-Par^5^ and commercial solutions like Antha^6^ and Transcriptics^7^. Graphical User Interfaces (GUIs) are commonly used to describe experiments but they are solution specific and do not allow interoperability.

### What are BioBlocks?

We developed BioBlocks to circumvent the programming bottleneck and allow users easier access to automation. BioBlocks is based on visual development environments like Google’s Blockly^8^ and Scratch^9^. They use a customizable toolbox of jigsaw-like blocks which can be linked together to produce a machine readable code (JSON, Python, etc.). BioBlocks has customized the blocks and grammar of Blockly, to allow the description of experimental protocols, in a simple drag and drop manner. The logic of BioBlocks is largely based on Autoprotocol^10^ (Supplementary Table 1), a language developed by Transcriptic for specifying experimental protocols in biology.

BioBlocks can be categorized into three types of blocks: ‘container blocks’, ‘operation blocks’ and ‘organization blocks’. Container blocks represent commonly used containers like multi-well plates, tubes, etc. ‘Operation blocks’ contain common procedures (actions) carried out during experimentation like pipetting, measuring, etc. Organization blocks help the user specify protocols in an intuitive manner akin to writing protocols in lab notebooks i.e. using Steps 1, Step 2, etc. Due care has been taken to ensure that visual manipulation of large protocols is easy; blocks or group of blocks can be minimized to allow for easy navigation between different parts of a protocol. These three types of blocks along with native Blockly blocks can be linked together iteratively to form large complex protocols (Figure 1). They can be saved, retrieved, modified and shared between multiple users.

**Figure 1.**
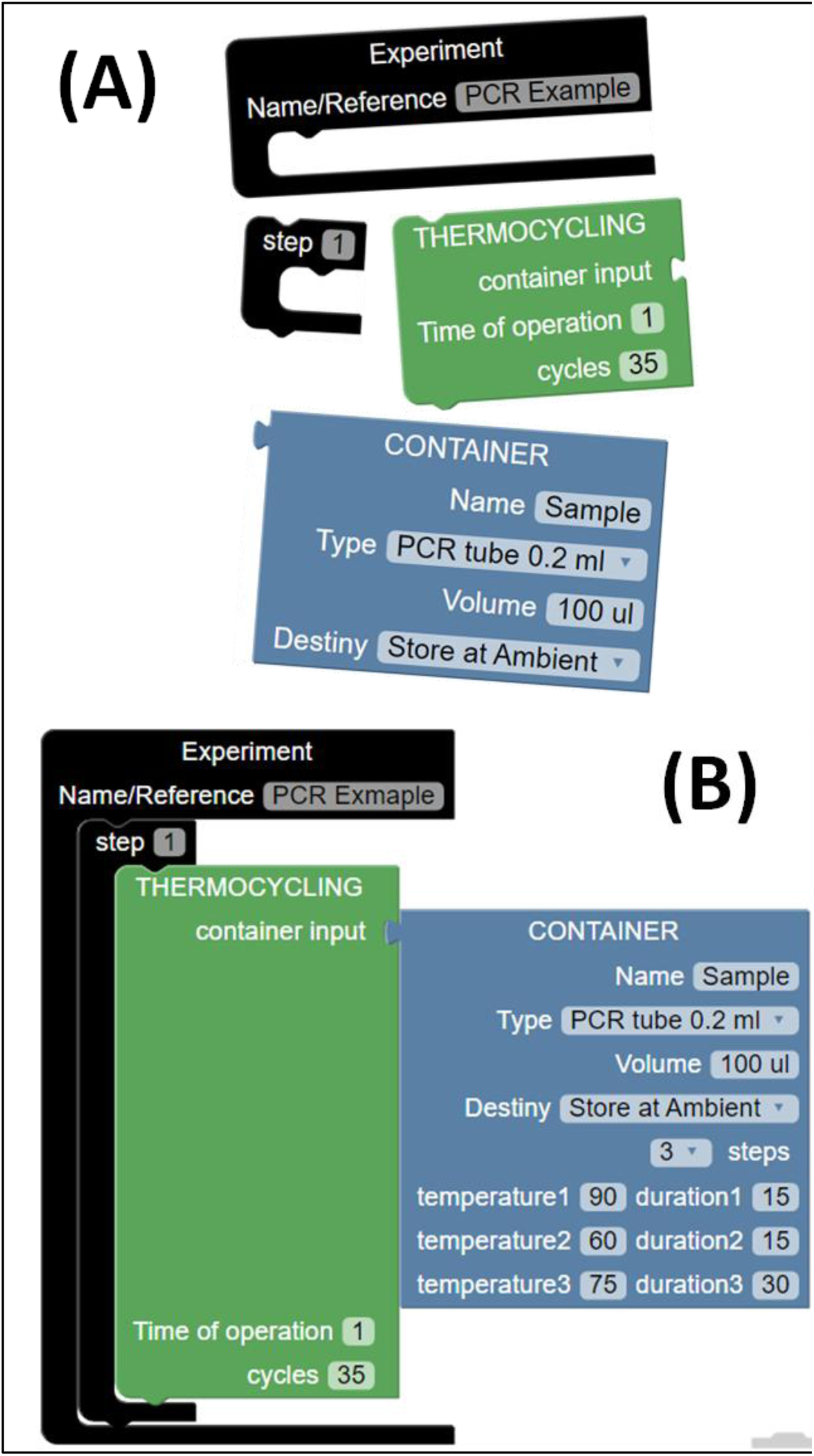
BioBlocks: A) Operation (green), Container (blue) and Organization (black) blocks needed to describe a PCR protocol are shown. B) The Blocks from A are linked together in a drag and drop way to describe a PCR protocol. The complimentary shapes of the blocks guide the user to build the protocols in an intuitive manner.

### Customization of Blocks

The open-source nature of Blockly gives user complete control over customization of the blocks. Customization can be done on two levels. First, modification of the blocks to generate machine code compatible for their choice of robotic platforms and execution of their protocols. Second, to introduce constraints to prohibit the linking of two incompatible blocks (i.e. the blocks snap away). The use of constraints in the design of the blocks helps avoid syntactic and logical errors. Syntactic errors are avoided because the code is generated in an automated manner and also due to the customization of blocks to the experimental biology domain. E.g. Operations like thermocycling are compatible only with specific types of containers (tubes). Applying this constraint restricts the user from linking incompatible containers like multi-well plates to a thermocycling block. Logical errors like overdrawing and under drawing fluid volumes can also be avoided. These constraints are encoded system-wide in the blocks. The user can create new blocks with different functionalities with a novel set of constraints or reuse/modify the existing constraints.

### BioBlocks Output

The protocols specified using BioBlocks are automatically translated in real-time to simultaneously generate multiple outputs (see Figure 2). The first output is a translation of the protocol to machine-readable code for its automated execution on a compatible hardware platform. The second output is a natural language (English) translation of the protocol to aid in verification. It is in the conventional format consisting of step-wise description of the protocol. The last output is the representation of the protocol as a workflow using Cytoscape. It is a powerful open-source tool which allows data analysis and visualization.^11^ The protocol workflow is an annotated and dynamic visualization, where the nodes and edges represent the containers and the action performed over the containers respectively. The goal of the workflow is to provide the user insight into planning and executing the protocol (Figure 2).

**Figure 2.**
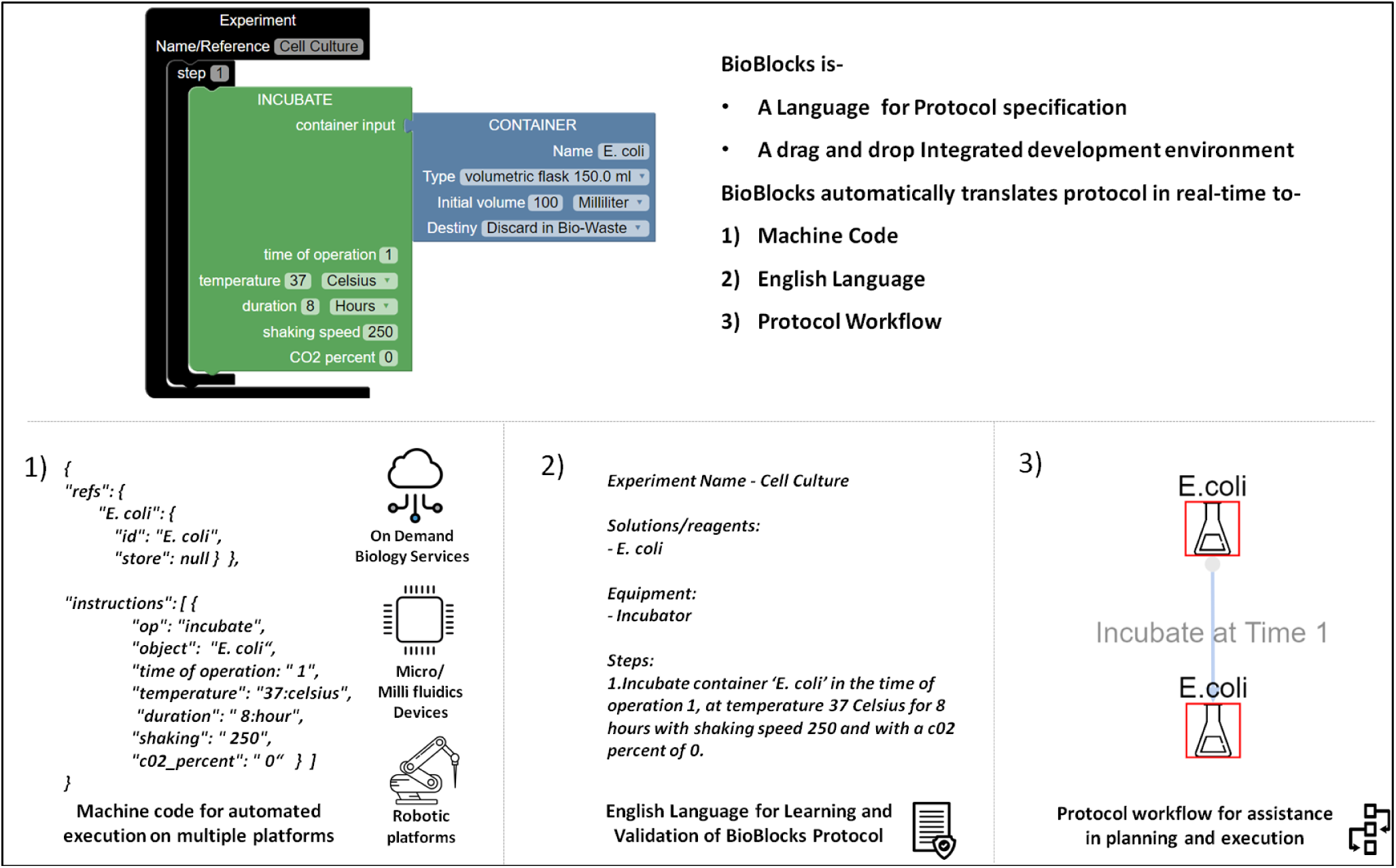
Multiple outputs of BioBlocks- An example of incubation of a cell culture in BioBlocks is shown. It is automatically translated to machine code (left), English (middle) and protocol workflow (right) in real time. The machine compatible code enables the user (non-programmer) to use automation and connect with multiple automation solutions.

As a proof of principle, the output code is generated using JSON syntax in two modes. The first mode is compatible with Autoprotocol, potentially allowing for remote execution of the described protocols at Transcriptics (a lab-in-a-cloud company). The second mode is an extension of Autoprotocol, which allows the description of protocols requiring feedback during execution e.g. continuous culture devices like turbidostats (Supplementary information) or any other new lab operation (block) included by the user. The real-time feedback and control of experiments, enables the user to guide the experiment based on real-time data. There are other drag and drop editors based on Autoprotocol (Wet Lab accelerator by Autodesk), but they do not allow conditional programming^12^ nor addition of new lab operations. As the DIY community for making open and 3D printable lab machines grows, BioBlocks would be very helpful for biologists to use it to operate those machines. ^13,14^

## CONCLUSION

We present BioBlocks, a web-based visual programming environment that addresses the problem of reproducibility by reducing ambiguity and minimizing human error using automation. On the front end, it is a visual programming interface based on the jigsaw model that has proven to be useful in multiple contexts. On the back end, the software system generates code compatible with lab automation settings for rapid prototyping of biological experiments. This work, is a step towards allowing the biologists to automate and codify their experiments in a simple manner, enabling to them to connect to multiple automation solutions.

## ACKNOWLEDGMENTS

This work was partially funded by EU FP7–FET-Proactive Project 610730 EVOPROG grant, Spanish National Projects TIN2012-36992 and TIN2016-81079-R grants. Icons in Figure 2 made by Freepik from www.flaticon.com

## SOFTWARE

Software, Tutorials and Code available on our webpage-http://www.lia.upm.es/index.php/software/Bioblocks

